# Influence of interindividual variability structures on implied priors and population predictions in pharmacogenetic models

**DOI:** 10.64898/2026.11.27.769637

**Authors:** Muhammad Waqar Ashraf, Nieko Punt, Laura Ramsey, Mikko Niemi, Klaus T. Olkkola, Vesa Kontinen, Teijo I. Saari

## Abstract

Interindividual variability (IIV) defines part of the prior distribution used in Bayesian parameter estimation. However, the dependence of model-implied priors on IIV and covariate model specifications is rarely examined explicitly. We evaluated empirical support for pooled versus phenotype-specific IIV structures and their consequences for predictive distributions in a CYP2D6-dependent population pharmacokinetic model. Four variants of a published codeine–metabolite model were independently re-estimated in NONMEM using data from 997 patients, combining continuous activity-score or categorical CYP2D6 parameterizations with pooled or phenotype-specific IIV. Distributions of morphine formation clearance were simulated in mrgsolve with 10,000 virtual individuals per phenotype and model. Two controlled 2 × 2 simulations isolated the effects of IIV variance structure within each parameterization. Phenotype-specific IIV improved model fit under both parameterizations (ΔOFV -448.57 and -44.27, respectively). The corresponding IIV variance estimates for poor metabolizers were 5.79 and 0.355. ETA variance shrinkage exceeded 90% in poor and ultrarapid metabolizers under both phenotype-specific IIV models. The four fitted models differed in the central tendency and dispersion of predicted morphine formation clearance. In controlled simulations, phenotype-specific IIV increased the coefficient of variation of morphine formation clearance in poor metabolizers approximately threefold under the continuous parameterization and by approximately 40% under the categorical parameterization. Both IIV estimates and their effects on population predictions depended on CYP2D6 parameterization. Importantly, improved model fit alone does not establish the biological validity of these distributions, and their implications for individual Bayesian posterior predictions remain to be evaluated.

## INTRODUCTION

For CYP2D6 substrates, genetic polymorphism is a major determinant of interindividual variability in drug metabolism. Pharmacokinetic studies have demonstrated substantial differences in drug disposition and exposure across CYP2D6 genotypes and phenotypes (Poulsen et al., 1996; Neafsey et al., 2009). CYP2D6-mediated metabolism is also substrate- and variant-dependent, as demonstrated by differences in the functional consequences of CYP2D6 variants (Johansson et al., 2025).

The specification of interindividual variability affects the covariate-conditional population distributions of individual pharmacokinetic parameters. These distributions serve as model-implied priors in Bayesian estimation, so the choice of variability structure may affect how strongly individual concentration data influence posterior estimates (Thomson and Whiting, 1992; Le Louedec et al., 2021). Pharmacogenetic effects on typical parameter values are commonly included as covariates, whereas interindividual variability is often assumed to be the same across phenotype groups. Dorne et al. (2002) found greater variability in oral clearance among non-phenotyped individuals and extensive metabolizers than among intermediate and poor metabolizers. Chiba et al. (2012) estimated genotype-specific coefficients of variation of 43–125% for CYP2D6 hepatic intrinsic clearance, while Darney et al. (2021) reported differences in pharmacokinetic variability across CYP2D6 phenotypes and populations. These findings suggest that a pooled interindividual variability (IIV) variance may not adequately describe variability within all phenotype groups.

Estimation of separate IIV variances for each phenotype increases model complexity and may yield imprecise estimates despite improved model fit, particularly when subgroup sizes are small or pharmacokinetic data are sparse. Tod et al. (2011) used Bayesian regression to estimate the contribution of CYP2D6-mediated metabolism to drug clearance and predict the magnitude of CYP2D6-mediated drug interactions. Grzegorzewski et al. (2022) incorporated variability in CYP2D6 enzyme kinetics derived from human liver microsomes into a PBPK model of dextromethorphan and examined variability even within individual CYP2D6 activity score groups. These approaches address CYP2D6-dependent parameter estimation and pharmacokinetic variability, but do not establish whether phenotype-specific IIV variances are supported by clinical data or how alternative variance structures affect simulated parameter distributions.

The aim of this study was to examine whether clinical data support phenotype-specific interindividual variability in the CYP2D6-mediated formation of morphine. We used the previously published population pharmacokinetic model of codeine and its metabolites (Ashraf et al., 2024). We compared four independently estimated models. The models combined continuous activity-score or categorical CYP2D6 fixed-effect parameterizations with pooled or phenotype-specific IIV variances. This comparison examined how model fit and estimated IIV variances depended on the specification of the CYP2D6 effect. Simulations from these models characterized the overall consequences of model specification for the predicted distributions of morphine formation clearance. In two separate controlled 2 × 2 simulation analyses, we applied both IIV variance structures to each fitted parameter set within the continuous and categorical CYP2D6 parameterizations, holding all other parameter estimates fixed. These comparisons isolated the effect of the variance structure within each fitted parameter set.

## METHODS

### Patient data

This secondary analysis used data from the CodAmpPain study (Poikola et al., 2025; Ashraf et al., 2024). The study protocol was approved by the Coordinating Ethics Committee of the HUS Hospital District (3/2016) and by the Finnish Medicines Agency Fimea (EudraCT number: 2015-005561-23). The study was registered prior to patient enrolment at the EU Clinical Trials Register (EudraCT 2015-005561-23). Written informed consent was obtained from all patients prior to participation in the study. The data was collected at the Day-Surgery Department of Jorvi Hospital, Helsinki University Hospital, Finland, during 8/2016–3/2018.

Patients received oral paracetamol (1,000 mg) and codeine (60 mg) as premedication. Plasma samples were collected approximately 20–60 and 180–360 min after administration, with an additional blood sample obtained for CYP2D6 genotyping. The clinical study, analytical methods, and genotyping procedures have been described previously (Poikola et al., 2025; Ashraf et al., 2024).

CYP2D6 activity scores and metabolizer phenotypes were assigned as described previously (Ashraf et al., 2024), following established genotype-to-phenotype translation recommendations (Gaedigk et al., 2008; Caudle et al., 2020; Crews et al., 2021). Scores of 0, greater than 0 but below 1.25, 1.25–2.25, and greater than 2.25 were defined poor, intermediate, normal, and ultrarapid metabolizers, respectively. In the original study, decreased-function alleles were assigned activity scores of 0.5, except for *CYP2D6*10*, which was assigned 0.25.

### Pharmacokinetic models

Of the 1,000 patients recruited for ambulatory surgery, 997 were included in the population pharmacokinetic analysis. The analyses were based on the previously published population pharmacokinetic model of codeine, morphine, codeine-6-glucuronide, and morphine-3-glucuronide (Ashraf et al., 2024). In the continuous activity-score parameterization, the model included CYP2D6 activity score (AS) as a continuous covariate on the fraction of codeine clearance directed towards morphine formation (*f*_MOR_). This fraction was parameterized on the logit scale, with AS centered at 2:

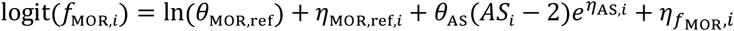

Here, *θ*_MOR,reF_ represents the typical population value of the morphine formation parameter at AS = 2, *θ*_AS_ describes the effect of AS on the logit scale, and 5_MOR,reF,i_ and 5_AS,i_ describe interindividual variability in these parameters. The additional random effect 5_ƒMOR,i_ represents the additional variability component examined in this study. The logistic transformation constrains *f*_MOR_ to the interval between 0 and 1.

In the alternative categorical parameterization, CYP2D6 metabolizer phenotype replaced continuous AS as the fixed-effect covariate. Separate parameters were estimated for poor (PM), intermediate (IM), normal (NM), and ultrarapid (UM) metabolizers:

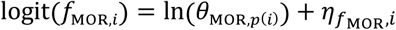

Here, *θ*_MOR,p_(_i_) represents the phenotype-specific parameter for individual *i*. The logistic transformation and the random effect on the logit scale were retained, whereas the additional random effects on the reference parameter and AS effect were absent from the categorical models.

Morphine formation clearance (*CL*_Form_) was calculated as the product of *f*_MOR_ and codeine clearance (*CL*_COD_):

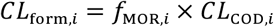

Morphine formation clearance was used to characterize the CYP2D6-dependent pathway in the simulation analyses.

### NONMEM re-estimation

Four variants of the population pharmacokinetic model were independently re-estimated using the original clinical data. The models combined two CYP2D6 fixed-effect parameterizations (continuous activity score and categorical metabolizer phenotype) with two IIV structures for the random effect on the logit of *f*_MOR_. The pooled-IIV models assumed a common variance across CYP2D6 phenotypes, whereas the phenotype-specific-IIV models estimated separate variances for PM, IM, NM, and UM. All four models retained the same structural and residual error models. Fixed-effect parameters and other estimable variance parameters were independently re-estimated in each model.

Parameters were estimated using the first-order conditional estimation method with interaction (FOCE-I) in NONMEM (version 7.6.0; ICON Development Solutions, Ellicott City, MD, USA), assisted by Perl-speaks-NONMEM (PsN) (Beal et al., 2026; Lindbom et al., 2005). Model fit was compared using the objective function value (OFV) and Akaike information criterion (AIC), calculated as OFV + 2k, where k is the number of estimated parameters. Comparisons between pooled and phenotype-specific IIV structures were made within each CYP2D6 fixed-effect parameterization. Comparisons between the continuous and categorical CYP2D6 parameterizations were treated as non-nested model comparisons. Parameter estimates, standard errors, relative standard errors (RSEs), estimated IIV variances, ETA shrinkage, and parameter correlations were examined to assess the estimation results.

### Model implementation and validation

The four independently re-estimated NONMEM models were implemented separately in mrgsolve (version 2.0.1; Elmokadem et al., 2019) using their respective final parameter estimates. The structural equations, parameter transformations, covariate relationships, and random-effects and residual error models were retained. All four mrgsolve implementations were validated against NONMEM predictions. Deterministic validation was performed with interindividual and residual variability disabled. Population predictions from NONMEM and mrgsolve were compared at 7,902 observation records per model across codeine, morphine, codeine-6-glucuronide, and morphine-3-glucuronide. Agreement was assessed using identity plots and prediction differences over time.

Stochastic validation assessed the implementation of interindividual and residual variability by comparing simulated random-effect and residual-error distributions with their specified distributions. Validation procedures and results are provided in the Supplementary Material.

### Simulation design and IIV parameterizations

A cohort of virtual individuals was generated by resampling patient covariates with replacement from the original dataset. The same phenotype-specific covariate cohorts were used to simulate all four independently fitted models, with 10,000 virtual individuals per CYP2D6 phenotype and model. Each model was simulated using its respective estimated parameters to characterize the distributions of the morphine formation fraction *f*_MOR_ and morphine formation clearance *CL*_Form_.

In a separate controlled 2 × 2 simulation using the continuous AS parameterization, we compared pooled and phenotype-specific IIV variance structures for *f*_MOR_ under both sets of fitted AS-model parameters, with all other parameters held fixed within each comparison. The two independently fitted models were supplemented with two hybrid simulation models, in which the IIV variance structure for *f*_MOR_ was exchanged while all other parameter estimates were retained. The hybrid models were used for simulation but were not re-estimated from the clinical data. For this analysis, 10,000 virtual individuals were simulated per phenotype and scenario, using the same resampled covariate cohorts across all four scenarios.

The same controlled 2 × 2 simulation design was applied to the categorical CYP2D6 parameterization. The two independently fitted categorical models were supplemented with two hybrid simulation models, in which only the *f*_MOR_ IIV variance structure was exchanged between the fitted parameter sets. All other fixed-effect parameters, random-effect variances, and residual error parameters were retained from the corresponding fitted model. Each scenario included 10,000 virtual individuals per phenotype, using identical resampled covariate cohorts but independent Monte Carlo random draws across scenarios.

To examine the implications for morphine exposure, plasma morphine concentration–time profiles were simulated using the two independently fitted continuous AS models. The same resampled covariate cohort of 10,000 virtual individuals per phenotype was used for both models, with independent draws of random effects. Individual plasma concentrations without residual error were simulated at 0.1-h intervals over 8 h following a single oral dose of 60 mg codeine. Median concentrations and empirical 5th–95th percentiles were calculated at each time point. Individual profiles were additionally visualized for 200 randomly selected individuals per phenotype, using the same selected individuals in both models.

Simulated *CL*_Form_ distributions were summarized using the mean, standard deviation (SD), median, 5th and 95th percentiles, and coefficient of variation (CV). Within each controlled 2 × 2 analysis, pooled and phenotype-specific IIV structures were compared separately under each set of fitted model parameters using ratios of distribution summary statistics and absolute differences in CV expressed in percentage points. These comparisons were based on independently simulated distributions rather than paired individual outcomes. Simulations used point estimates of model parameters without propagating parameter estimation uncertainty. Data processing, simulations, and graphical analyses were performed using R (version 4.5.2; R Core Team, 2025) with RStudio (Posit Software; Posit Team, 2025, https://posit.co/products/open-source/rstudio) and mrgsolve (version 2.0.1).

### Data and Code Availability

The individual-level clinical data used in this study are not publicly available due to restrictions related to participant confidentiality and the conditions governing the original clinical study. The NONMEM model specifications, mrgsolve implementations, parameter estimates, simulation scripts, and a synthetic example dataset are available in a public repository [https://doi.org/10.5281/zenodo.24510573]. The example dataset is intended to demonstrate the computational workflow and does not reproduce the original clinical data. Exact reproduction of analyses involving the observed patient covariates or clinical concentration measurements requires access to the restricted source dataset.

## RESULTS

Of the 1,000 patients recruited, 997 had the required plasma concentration and CYP2D6 genotype data for population pharmacokinetic analysis. A total of 64 distinct CYP2D6 genotypes were identified and classified by activity score into poor (PM, n = 37), intermediate (IM, n = 268), normal (NM, n = 629), and ultrarapid (UM, n = 63) metabolizer phenotypes.

### Population pharmacokinetic model estimation and variability parameterization

The phenotype-specific IIV models provided a better fit than the corresponding pooled-IIV models under both CYP2D6 fixed-effect parameterizations (Table 1). With continuous AS, the OFV decreased from −1911.394 to −2359.967 (ΔOFV = −448.573), whereas with categorical phenotypes, it decreased from −2372.330 to −2416.598 (ΔOFV = −44.268). Each comparison involved three additional variance parameters. The corresponding AIC values were −1869.394 and −2311.967 for the continuous AS models and −2326.330 and −2364.598 for the categorical models, respectively. The categorical parameterization yielded lower OFV and AIC values than the continuous AS parameterization under both IIV structures.

**Table 1.** Comparison of CYP2D6 fixed-effect parameterizations and interindividual variability structures.

| **Characteristic** | **Activity score** |  | **Categorical** |  |
|---|---|---|---|---|
|  | Pooled IIV | Grouped IIV | Pooled IIV | Grouped IIV |
| Number of estimated parameters (k) | 21 | 24 | 23 | 26 |
| Objective function value (OFV) | -1911.394 | -2359.967 | -2372.330 | -2416.598 |
| Akaike information criterion (AIC) | -1869.394 | -2311.967 | -2326.330 | -2364.598 |
| ΔOFV vs pooled IIV | - | -448.573 | - | -44.268 |
| **IIV variance of fMOR (ω²)** |  |  |  |  |
| Common (pooled) | 0.3338 | - | 0.1848 | - |
| PM | - | 5.7952 | - | 0.3554 |
| IM | - | 0.2341 | - | 0.2891 |
| NM | - | 0.1189 | - | 0.1329 |
| UM | - | 0.3262 | - | 0.1100 |

Values are from independently re-estimated NONMEM models fitted to the same dataset (997 individuals; 7,902 concentration observations). Fixed-effect CYP2D6 parameterizations were either continuous activity score (AS) or four categorical metabolizer phenotypes. Grouped IIV denotes separate variances for PM, IM, NM, and UM; pooled IIV denotes one common variance.
ΔOFV is the difference between grouped and pooled models within the same CYP2D6 fixed-effect parameterization (three additional estimated variance parameters in each comparison). IIV variances (ω²) refer to the random effect on the log-odds of the fraction of codeine clearance directed to morphine formation (*f*<sub>*MOR*</sub>). AIC = OFV + 2k.
Abbreviations: IIV, interindividual variability; PM, poor metabolizer; IM, intermediate metabolizer; NM, normal metabolizer; UM, ultrarapid metabolizer.

Re-estimation also changed fixed-effect estimates, including the reference population value of the morphine formation parameter (0.1584 vs. 0.09522) and the CYP2D6 activity score effect on the logit scale (1.0042 vs. 0.7119) for the pooled and phenotype-specific models, respectively (Supplementary Table S1). The estimated *f*_MOR_ variance was 0.334 in the pooled model, compared with 5.79, 0.234, 0.119, and 0.326 for PM, IM, NM, and UM, respectively, in the phenotype-specific model (Supplementary Table S2). The largest difference between parameterizations was observed in PM, where the phenotype-specific variance was substantially higher with continuous AS than with categorical phenotypes (Table 1; Supplementary Tables S2 and S3).

In the phenotype-specific IIV models, ETA variance shrinkage was particularly high for PM (96.3% and 96.9%) and UM (94.7% and 96.0%) under the continuous AS and categorical parameterizations, respectively. In the pooled-IIV models, overall ETA variance shrinkage was 16.6% and 25.1%, respectively. These overall estimates are not directly comparable with the phenotype-specific shrinkage estimates. The relative standard errors of the random-effect standard deviation estimates were 30.0% and 16.1% for PM and 10.0% and 10.8% for UM, respectively (Supplementary Tables S2 and S3). The categorical pooled-IIV model showed relative standard errors exceeding 100% for several structural parameters, accompanied by strong correlations among selected fixed-effect parameters (Supplementary Table S3). Estimation of an exploratory model combining selected CYP2D6 phenotype groups terminated with numerical errors, and the covariance step failed (Supplementary Table S4).

### Model translation and validation

The four independently re-estimated population pharmacokinetic models were successfully implemented in mrgsolve. Minimization and covariance steps were completed for all four models, although NONMEM reported a minimization warning for the continuous AS pooled-IIV model (Supplementary Table S3). Deterministic validation showed close agreement between NONMEM and mrgsolve population predictions across all four analytes, with 7,902 observation records evaluated for each model. The maximum absolute differences on the log-concentration scale were 0.00010 and 0.00077 for the continuous AS pooled- and phenotype-specific-IIV models, respectively, and 0.000054 and 0.000054 for the corresponding categorical models. Stochastic validation confirmed the implementation of the random-effects and residual error structures in all four models. Simulated residual errors were centered at zero, with analyte-specific dispersion consistent with the specified residual error model. Deterministic validation of the original published model is illustrated in Supplementary Figures S1–S2, and stochastic residual error validation of the two continuous AS models in Supplementary Figure S3.

### Model-implied distributions of morphine formation clearance

The four independently fitted models produced different distributions of morphine formation clearance across CYP2D6 phenotypes (Figure 1). Each model was simulated using its own estimated parameter set, with 10,000 virtual individuals per phenotype. Differences were observed in both the central tendency and dispersion of morphine formation clearance, depending on the CYP2D6 fixed-effect parameterization and IIV structure. The largest differences were observed in poor metabolizers.

**Figure 1.**
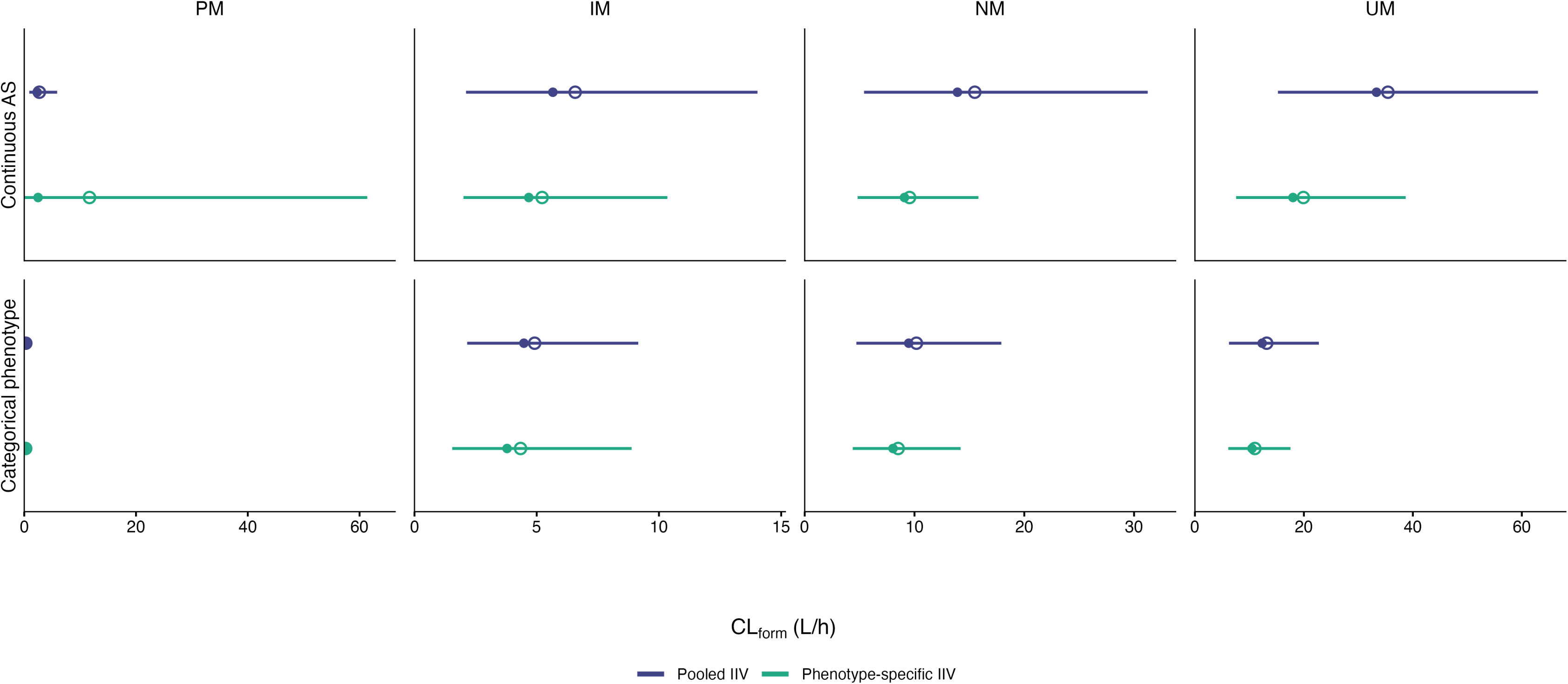
Model-implied distributions of morphine formation clearance across four independently fitted pharmacokinetic models. Median (filled circles), mean (open circles), and empirical 5th–95th percentiles (horizontal lines) are shown for poor (PM), intermediate (IM), normal (NM), and ultrarapid (UM) metabolizers. The upper and lower rows represent the continuous activity-score and categorical CYP2D6 parameterizations, respectively. Blue and green denote pooled and phenotype-specific IIV structures. Each model was simulated using its respective estimated parameter set and the same resampled covariate cohort of 10,000 virtual individuals per phenotype. The horizontal axes are linear, with phenotype-specific scales. IIV, interindividual variability.

Under phenotype-specific IIV, the continuous AS model produced a median morphine formation clearance of 2.48 L/h and a coefficient of variation of 178.2%, compared with 0.215 L/h and 66.3%, respectively, under the categorical parameterization. The distributions therefore differed substantially in both location and dispersion. Differences between the fitted models were also observed in intermediate, normal, and ultrarapid metabolizers, including differences in median morphine formation clearance among ultrarapid metabolizers (Figure 1; Supplementary Table S5). The distributions for the two continuous AS models are additionally illustrated as violin plots in Supplementary Figure S4. Because all four models were independently re-estimated, these differences reflect changes in the estimated fixed effects and other model parameters as well as the IIV structures.

### Controlled effects of IIV variance structures on morphine formation clearance

We performed a controlled 2 × 2 simulation analysis using the continuous AS parameterization to examine the effect of IIV variance structure independently of other differences between the two fitted models. Pooled and phenotype-specific IIV structures were evaluated under both sets of estimated model parameters, with all other parameters held constant within each comparison. The four scenarios included 10,000 simulated individuals per CYP2D6 phenotype, using identical resampled covariate cohorts across scenarios (Table 2, Figure 2).

**Figure 2.**
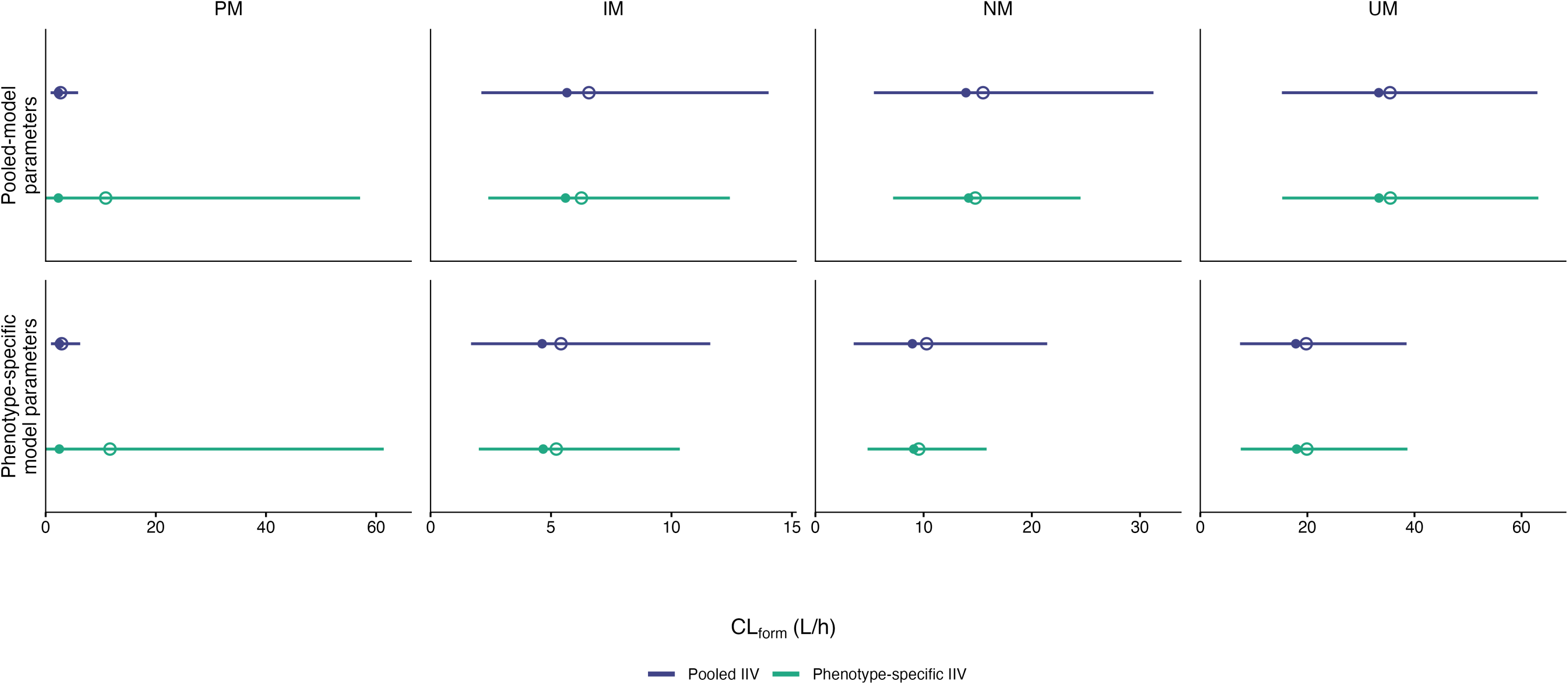
Effect of phenotype-specific variability on simulated morphine formation clearance under the continuous activity-score parameterization. Median (filled circles), mean (open circles), and 5th–95th percentile intervals (horizontal lines) of simulated morphine formation clearance are shown for poor (PM), intermediate (IM), normal (NM), and ultrarapid (UM) metabolizers. The upper row uses parameter estimates from the pooled model, whereas the lower row uses those from the phenotype-specific model. Within each row, only the IIV variance structure is varied, with all other model parameters held constant. Each scenario includes 10,000 virtual individuals per phenotype, using identical resampled covariate cohorts across scenarios. The horizontal axis is linear, with phenotype-specific scales. The two hybrid scenarios were used for simulation only and were not fitted to the clinical data. IIV, interindividual variability.

**Table 2.** Controlled 2 × 2 simulation analysis of morphine formation clearance by CYP2D6 phenotype. Each scenario comprised 10,000 simulated individuals per phenotype, with the same resampled covariate cohorts. The random effects were drawn independently between scenarios.

| **Phenotype** | **Scenario** | **Mean (SD)** | **Median** | **5th–95th percentile** | **CV (%)** |
|---|---|---|---|---|---|
| **PM** | A | 2.70 (1.66) | 2.31 | 0.89–5.90 | 61.38 |
|  | B | 10.90 (19.82) | 2.31 | 0.04–57.09 | 181.82 |
|  | C | 2.90 (1.75) | 2.48 | 0.96–6.27 | 60.35 |
|  | D | 11.68 (20.81) | 2.48 | 0.05–61.38 | 178.23 |
| **IM** | A | 6.57 (3.94) | 5.66 | 2.11–14.03 | 59.90 |
|  | B | 6.26 (3.20) | 5.60 | 2.40–12.42 | 51.03 |
|  | C | 5.41 (3.31) | 4.63 | 1.68–11.61 | 61.16 |
|  | D | 5.22 (2.69) | 4.68 | 2.00–10.35 | 51.46 |
| **NM** | A | 15.49 (8.14) | 13.91 | 5.40–31.25 | 52.54 |
|  | B | 14.77 (5.37) | 14.16 | 7.18–24.51 | 36.35 |
|  | C | 10.28 (5.82) | 8.96 | 3.54–21.42 | 56.63 |
|  | D | 9.55 (3.44) | 9.09 | 4.82–15.82 | 35.97 |
| **UM** | A | 35.42 (14.75) | 33.33 | 15.24–62.96 | 41.63 |
|  | B | 35.49 (14.76) | 33.37 | 15.29–63.14 | 41.58 |
|  | C | 19.78 (9.95) | 17.84 | 7.42–38.53 | 50.32 |
|  | D | 19.91 (9.99) | 17.99 | 7.56–38.69 | 50.17 |

Morphine formation clearance is expressed in L/h. Scenarios A and B use the pooled-model parameter estimates, while scenarios C and D use the phenotype-specific model parameter estimates. Scenarios A and C use a pooled variance for the random effect governing morphine formation; B and D use phenotype-specific variances.  Within each parameter set, only the *f*<sub>MOR</sub>  IIV variance structure changes. B and C are hybrid simulation scenarios, not independently estimated models.  PM, poor metabolizer; IM, intermediate metabolizer; NM, normal metabolizer; UM, ultrarapid metabolizer; IIV, interindividual variability; CV, coefficient of variation.

Under the pooled-model parameter estimates, replacing pooled with phenotype-specific *f*_MOR_ increased mean (SD) morphine formation clearance in poor metabolizers from 2.70 (1.66) to 10.90 (19.82) L/h and the coefficient of variation from 61.38% to 181.82%. Under the phenotype-specific model parameter estimates, the corresponding mean (SD) increased from 2.90 (1.75) to 11.68 (20.81) L/h, with the coefficient of variation increasing from 60.35% to 178.23%. Despite these substantial changes, median formation clearance remained unchanged at 2.31 L/h under the pooled-model parameter estimates and at 2.48 L/h under the phenotype-specific model parameter estimates, regardless of the IIV variance structure applied. The 5th–95th percentile intervals widened from 0.89–5.90 to 0.04–57.09 L/h with the pooled-model parameters and from 0.96–6.27 to 0.05–61.38 L/h with the phenotype-specific model parameters (Table 2, Figure 2).

In intermediate metabolizers, phenotype-specific IIV reduced the coefficient of variation from 59.90% to 51.03% under the pooled-model parameters and from 61.16% to 51.46% under the phenotype-specific model parameters. The corresponding reductions in normal metabolizers were from 52.54% to 36.35% and from 56.63% to 35.97%, respectively. In ultrarapid metabolizers, the coefficient of variation remained nearly unchanged (41.63% vs. 41.58% and 50.32% vs. 50.17%, respectively) (Table 2).

The corresponding controlled analysis using categorical CYP2D6 phenotypes showed smaller changes in morphine formation clearance distributions among poor metabolizers. Replacing pooled with phenotype-specific IIV increased the coefficient of variation by 19.4 and 18.2 percentage points under the pooled- and phenotype-specific model parameter estimates, respectively, compared with approximately 120 and 118 percentage points under the continuous AS parameterization. In intermediate metabolizers, the corresponding increases were 10.6 and 11.2 percentage points, whereas the coefficient of variation decreased by 5.8 percentage points in normal metabolizers and 8.2 percentage points in ultrarapid metabolizers under both parameter sets. Median formation clearance remained nearly unchanged within each controlled comparison (Supplementary Table S6).

### Model-implied morphine concentration–time profiles

Differences between the two independently fitted continuous AS models were also reflected in simulated morphine concentration–time profiles. The 5th–95th percentile interval was markedly wider in poor metabolizers under the phenotype-specific IIV model, despite relatively similar median profiles (Figure 3). Differences between models were less pronounced in the other phenotypes. Individual simulated profiles further illustrated the differences in between-subject variability (Supplementary Figure S5).

**Figure 3.**
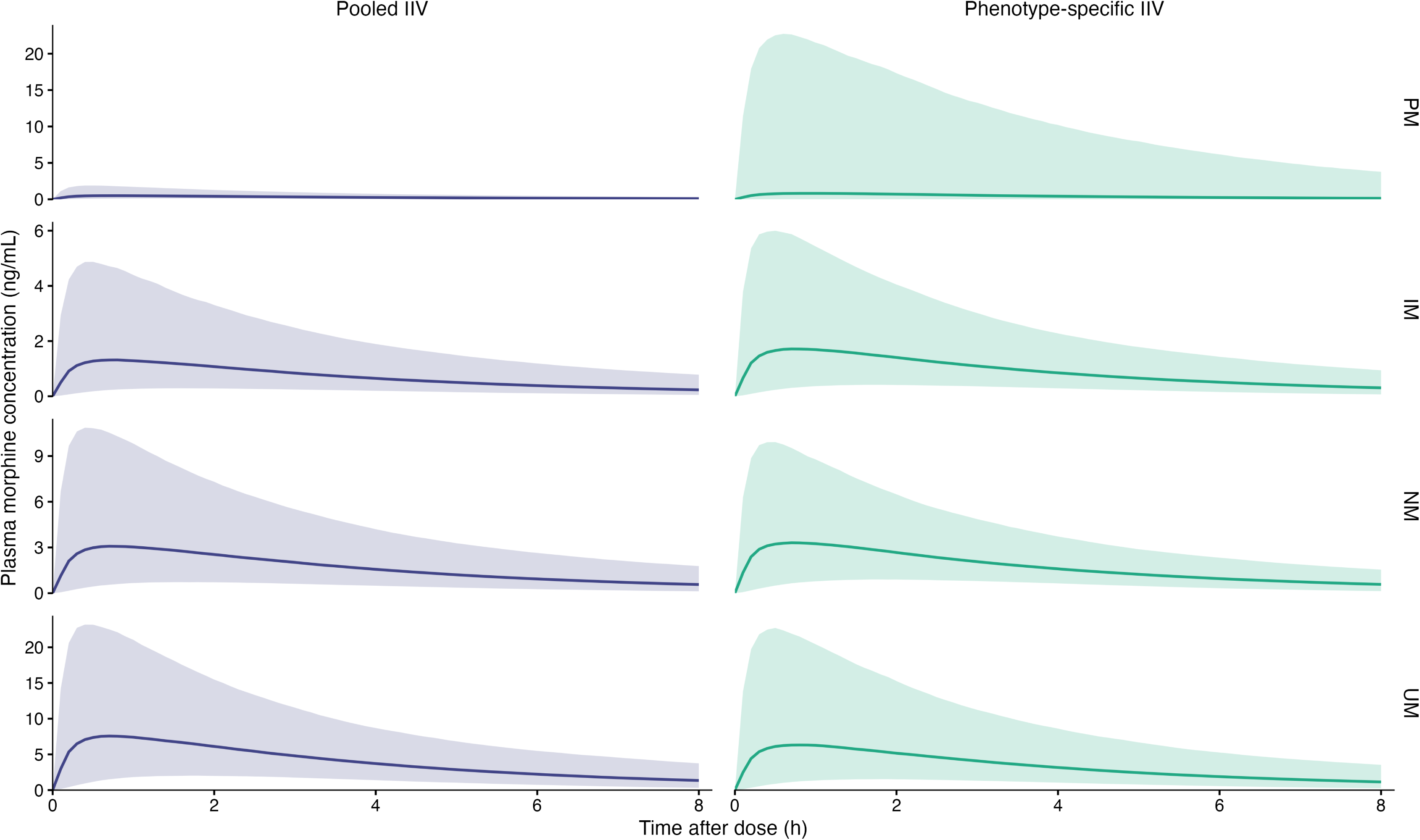
Simulated morphine concentration–time profiles by CYP2D6 phenotype under the continuous activity-score parameterization. Solid lines represent median simulated plasma morphine concentrations, and shaded areas indicate the empirical 5th–95th percentile intervals following a single oral dose of 60 mg codeine. Simulated concentrations do not include residual error. Each model was simulated using its own estimated parameter set and the same resampled cohort of 10,000 virtual individuals per phenotype. Results are shown for poor (PM), intermediate (IM), normal (NM), and ultrarapid (UM) metabolizers under the pooled (blue) and phenotype-specific (green) IIV models. Concentration axes differ between phenotypes but are identical between models within each phenotype.

## DISCUSSION

This study evaluated the empirical support for pooled and phenotype-specific IIV in a CYP2D6-dependent population pharmacokinetic model and examined the predictive distributions implied by the four fitted models. Phenotype-specific IIV improved model fit under both CYP2D6 parameterizations. However, the estimated IIV variances differed between parameterizations, particularly in poor metabolizers and individual random-effect estimates were poorly informed in some phenotypes. The four fitted models differed in both the central tendency and dispersion of predicted morphine formation clearance. The effects of changing the IIV variance structure also differed between CYP2D6 parameterizations in the controlled simulations. In poor metabolizers, phenotype-specific IIV increased the coefficient of variation of morphine formation clearance by approximately 120 percentage points under the continuous activity-score parameterization and by 18– 19 percentage points under the categorical parameterization. Thus, both the estimated IIV variances and the effects of alternative IIV structures on predicted parameter distributions depended on the CYP2D6 model specification.

Nyberg and Jonsson (2025) showed that covariate model misspecification can affect the estimation of IIV variances. In our study, the categorical parameterization yielded a smaller PM variance estimate with greater relative precision, but ETA shrinkage remained high. As discussed by Savic and Karlsson (2009), high ETA shrinkage reflects limited information in the individual data relative to the population distribution. In our analysis, improved precision of the population variance estimate was therefore not accompanied by reduced shrinkage of the individual random-effect estimates. The two CYP2D6 parameterizations also differed in their random-effects structures. The estimated variances therefore represent specific variance components rather than directly comparable measures of total within-phenotype variability. The uncertainty and correlations of fixed-effect parameter estimates suggest that neither parameterization fully resolved the problems of parameter identifiability. Thus, improved overall model fit does not necessarily imply that variability can be reliably characterized within every phenotype. These observations support the recommendations of Sanghavi et al. (2024) to consider parameter uncertainty when evaluating covariate models. High ETA shrinkage may also reflect the limited information provided by sparse concentration sampling and insufficient individual identifiability of morphine formation, rather than subgroup size alone.

The differences between the distributions of morphine formation clearance reflect the combined effects of fixed-effect estimates and IIV structures. Their biological interpretation requires consideration of previous clinical findings. Reduced morphine formation in poor metabolizers is well established (Crews et al., 2021). In a clinical study of codeine pharmacokinetics, neither morphine nor morphine-6-glucuronide was detectable in 13 of 14 poor metabolizers (Poulsen et al., 1996). The lower morphine formation clearance predicted by the categorical models in poor metabolizers appears more consistent with these clinical observations. However, the available clinical evidence does not establish the extent of within-phenotype variability in morphine formation clearance. Neither CYP2D6 parameterization is based on direct measurements of individual enzyme activity as both represent structural assumptions about genotype-dependent morphine formation. Improved model fit should therefore not be equated with greater biological or clinical validity.

Our controlled simulations showed that consequences of alternative IIV structures depend on the CYP2D6 parameterization. In poor metabolizers, mean formation clearance increased approximately fourfold without a corresponding change in the median under the continuous activity-score parameterization. The corresponding increase was smaller under the categorical parameterization. The mean of a logit-normal distribution depends on both the mean and variance of the underlying normal distribution (Holmes and Schofield, 2022). Thus, the fixed-effect estimates alone do not determine the expected morphine formation clearance within a phenotype.

Previous studies have demonstrated variability in CYP2D6-mediated metabolism both between and within metabolizer phenotypes (Dorne et al., 2002; Chiba et al., 2012; Darney et al., 2021). These observations support the consideration of phenotype-specific IIV in pharmacogenetic models.

Mechanistic PBPK studies have further examined variability within CYP2D6 activity score groups. Grzegorzewski et al. (2022) incorporated variability in CYP2D6 enzyme kinetics into their model, whereas Rüdesheim et al. (2022) used individual concentration profiles to characterize variability in CYP2D6 catalytic activity. Such mechanistic approaches complement population pharmacokinetic approaches by examining potential sources of within-phenotype variability. Furthermore, Rüdesheim et al. reported insufficient data to characterize variability in the lowest activity score groups. Our findings suggest similar difficulties in characterizing variability in poor metabolizers, irrespective of the CYP2D6 parameterization.

Previous work has demonstrated the value of prior information in quantitative modelling of CYP2D6-mediated drug metabolism (Tod et al., 2011). The relevance of our findings to Bayesian forecasting lies in the use of covariate-conditional population distributions as priors for individual parameter estimation. The simulated phenotype-specific distributions additionally incorporate within-phenotype covariate variability. In the context of Bayesian therapeutic drug monitoring, Baklouti et al. (2022) examined the consequences of shrinkage for individual pharmacokinetic parameter estimates. Wicha et al. (2021) emphasized the need for model validation before clinical application in model-informed precision dosing. Our findings suggest that the specification of both fixed effects and IIV warrants consideration in the selection of pharmacogenetic models for Bayesian forecasting. This may be particularly relevant when individual concentration data provide limited information about the parameter of interest. However, we did not evaluate the consequences of alternative model-implied priors for individual posterior predictions or clinical dosing decisions.

Several limitations should be acknowledged. All four models were based on the same clinical dataset and shared the same structural pharmacokinetic model. Sparse concentration sampling and unequal phenotype group sizes limited the information available for parameter estimation. The forward simulations used point estimates of model parameters without accounting for their estimation uncertainty. Uncertainty in the estimated codeine clearance also limits the interpretation of absolute morphine formation clearance values. The results of the controlled simulations were therefore conditional on the estimated parameter sets. The predictive performance of the alternative models was not evaluated in an independent clinical dataset. We did not evaluate individual Bayesian posterior predictions or clinical dosing outcomes.

In conclusion, phenotype-specific IIV improved likelihood-based model fit under both CYP2D6 parameterizations, but the estimated variances depended on the specification of the CYP2D6 effect. The four fitted models implied different distributions of morphine formation clearance. The controlled simulations showed that the effects of alternative IIV structures on these distributions also depended on the CYP2D6 parameterization. Both fixed-effect and IIV specifications should therefore be considered when evaluating the model-implied priors of pharmacogenetic models for Bayesian forecasting.

## Study Highlights

### What is the current knowledge on the topic?

**Pharmacogenetic** effects on drug disposition are well established, and population pharmacokinetic models commonly incorporate genotype or phenotype as covariates. Interindividual variability is typically pooled across subgroups for pragmatic reasons, but its role as an implicit prior in Bayesian prediction is rarely examined.

### What question did this study address?

Do clinical data support phenotype-specific variability in the CYP2D6-dependent morphine formation pathway, and how do the alternative fitted models differ in their implied population predictions?

### What does this study add to our knowledge?

Phenotype-specific *f*_MOR_ IIV improved likelihood-based fit but showed very high ETA shrinkage in the PM and UM groups. Independently fitted models differed in both central tendency and dispersion of simulated morphine formation clearance.

### How might this change drug discovery, development, and/or therapeutics?

Explicit evaluation of IIV assumptions may improve transparency of uncertainty in model-informed precision dosing, particularly in sparsely represented pharmacogenetic subgroups.

## Acknowledgements

The authors used ChatGPT (OpenAI) for editorial feedback and assistance with wording during manuscript preparation. The authors reviewed and edited the resulting text and take responsibility for the final manuscript.

## Author Contributions

All authors contributed to writing the manuscript. M.N., K.T.O., and V.K. designed the original clinical study. V.K. conducted the original clinical study. M.W.A., N.P., L.R. and T.I.S. designed the present analysis and analyzed the data. T.I.S. performed the simulations and led manuscript drafting and finalization.

## Supplementary Information Titles

**Supplementary Material.** Model estimation diagnostics, exploratory phenotype grouping, NONMEM-to-mrgsolve validation, and additional simulation results (Tables S1–S6 and Figures S1– S5)

