## Supplementary material for "Influence of interindividual variability structures on implied priors and population predictions in pharmacogenetic models"

### 1 Estimation diagnostics

The original clinical data from 997 patients were used to independently re-estimate four variants of the population pharmacokinetic model by first-order conditional estimation with interaction in NONMEM. The models combined two CYP2D6 fixed-effect parameterizations (continuous activity score and categorical metabolizer phenotype) with pooled or phenotype-specific interindividual variability (IIV) for the random effect governing the morphine formation fraction $f_{\mathrm{MOR}}$. All four models retained the same structural and residual error models, while fixed-effect parameters and estimable variance parameters were independently re-estimated.

Phenotype-specific IIV improved likelihood-based fit under both CYP2D6 parameterizations. With continuous activity score, the objective function value (OFV, without the constant) decreased from -1911.39 to -2359.97 ($\Delta$OFV $= -448.57$), whereas with categorical phenotypes it decreased from -2372.33 to -2416.60 ($\Delta$OFV $= -44.2683$). Each comparison involved three additional variance parameters. The corresponding Akaike information criterion values were -1869.39 and -2311.97 for the continuous activity-score models and -2326.33 and -2364.60 for the categorical models. These differences indicate improved likelihood-based fit but do not establish superior individual Bayesian posterior prediction. Fixed-effect parameter estimates are presented in Supplementary Table S1, the original continuous activity-score model IIV estimates and shrinkage diagnostics in Supplementary Table S2, and estimation precision, shrinkage, and selected parameter correlations across all four models in Supplementary Table S3.

### 2 Exploratory alternative phenotype grouping

The exploratory model combined poor and intermediate metabolizers into a single IIV variance group, while retaining separate variances for normal and ultrarapid metabolizers. Its original diagnostics reported rounding/termination problems and a failed covariance step; consequently, parameter RSEs were unavailable. The model was not used in the primary forward simulation comparison. Its OFV was not used in place of that of the primary continuous activity-score model with phenotype-specific IIV (Supplementary Table S4).

### 3 Model translation and validation

The previously published NONMEM population pharmacokinetic model served as the basis for four model variants combining two CYP2D6 fixed-effect parameterizations (continuous activity score and categorical phenotype) with pooled or phenotype-specific IIV structures. All four variants were independently re-estimated using the original clinical data and subsequently implemented in mrgsolve for simulation. The original published model was first translated into mrgsolve and evaluated using deterministic validation, with interindividual and residual variability disabled (Supplementary Figures S1–S2). The four re-estimated models were then validated separately against NONMEM predictions. Deterministic comparisons included 7,902 observation records per model. The maximum absolute differences in log-concentration predictions were $1.0 \times 10^{-4}$ and $7.7 \times 10^{-4}$ for the continuous activity-score pooled- and phenotype-specific-IIV models, respectively, and $5.4 \times 10^{-5}$ for both categorical models.

Stochastic validation evaluated simulated interindividual random effects and residual errors against their specified variance structures. For the continuous activity-score models, simulations included 3,000 virtual individuals per CYP2D6 phenotype for each model. All 31 $\Omega$- and eight $\Sigma$-variance checks met the prespecified acceptance criteria. For the categorical models, all 40 variance checks (eight $\Omega$- and 32 $\Sigma$-checks across both models) met the acceptance criterion of a relative deviation not exceeding 15%. The maximum absolute relative deviations were 6.52% and 4.23% for the categorical pooled- and phenotype-specific-IIV models, respectively. Supplementary Figure S3 illustrates the residual error distributions for the continuous activity-score models.

**Supplementary Table S1:** Fixed-effect parameter estimates across four CYP2D6 model specifications.

| | Continuous activity score | | Categorical phenotype | |
|---|---|---|---|---|
| **Parameter description** | **Pooled IIV** | **Grouped IIV** | **Pooled IIV** | **Grouped IIV** |
| Codeine absorption rate ($K_a$, $h^{-1}$) | 8.74 (13.2) | 8.43 (13.4) | 8.92 (12.8) | 8.87 (12.8) |
| Codeine clearance ($CL_{COD}$, $L\,h^{-1}$) | 110 (28.4) | 110 (40.6) | 103 (110.2) | 102 (44.9) |
| Codeine volume ($V_{COD}$, L) | 428 (28.6) | 420 (40.6) | 394 (110.2) | 391 (45.0) |
| Codeine bioavailability ($F$) | 0.839 (28.4) | 0.832 (40.5) | 0.781 (110.3) | 0.775 (45.0) |
| Morphine clearance ($CL_{MOR}$, $L\,h^{-1}$) | 358 (31.4) | 226 (47.2) | 229 (113.4) | 195 (49.4) |
| Morphine volume ($V_{MOR}$, L) | 22.8 (34.9) | 13.4 (50.2) | 14.09 (114.6) | 11.65 (51.3) |
| C6G clearance ($CL_{C6G}$, $L\,h^{-1}$) | 7.97 (28.6) | 8.38 (40.8) | 7.80 (110.3) | 7.84 (45.4) |
| C6G volume ($V_{C6G}$, L) | 5.37 (28.5) | 5.67 (40.9) | 5.33 (110.4) | 5.35 (45.5) |
| M3G clearance ($CL_{M3G}$, $L\,h^{-1}$) | 9.44 (31.4) | 5.96 (47.1) | 6.05 (113.4) | 5.15 (49.3) |
| M3G volume ($V_{M3G}$, L) | 8.47 (31.7) | 5.39 (47.3) | 5.50 (113.3) | 4.69 (49.4) |
| **CYP2D6 fixed-effect parameters** | | | | |
| Reference morphine formation parameter (AS = 2) | 0.158 (16.0) | 0.0952 (29.4) | — | — |
| AS effect on morphine formation | 1.00 (2.12) | 0.712 (4.74) | — | — |
| PM formation parameter | — | — | 0.00246 (25.8) | 0.00208 (31.2) |
| IM formation parameter | — | — | 0.0467 (26.4) | 0.0396 (31.3) |
| NM formation parameter | — | — | 0.1019 (26.7) | 0.0861 (31.4) |
| UM formation parameter | — | — | 0.1376 (27.7) | 0.1161 (31.9) |

Estimates are reported as point estimate (relative standard error, %). Dashes indicate parameters absent from the respective fixed-effect model. The categorical model uses four phenotype-specific formation parameters (PM, IM, NM, UM), whereas the continuous AS model uses a reference formation parameter and an AS effect. Parameter descriptions are aligned across models because the THETA indices differ between parameterizations. The fixed THETA(15) (fraction of morphine metabolism to M3G, 0.6) is not included.

**Supplementary Table S2:** $f_{\mathrm{MOR}}$ random-effect estimates and ETA shrinkage.

| Model / phenotype | ETA SD ($\omega$) | OMEGA variance ($\omega^2$) | RSE of ETA SD (%) | ETA shrinkage SD (%) | ETA shrinkage variance (%) |
|---|---|---|---|---|---|
| Pooled / all | 0.578 | 0.334 | 2.10 | 8.68 | 16.6 |
| Phenotype-specific / PM | 2.407 | 5.79 | 30.0 | 80.9 | 96.3 |
| Phenotype-specific / IM | 0.484 | 0.234 | 3.68 | 53.7 | 78.6 |
| Phenotype-specific / NM | 0.345 | 0.119 | 2.16 | 35.6 | 58.6 |
| Phenotype-specific / UM | 0.571 | 0.326 | 10.0 | 77.0 | 94.7 |

$\omega$ is the standard deviation of the $f_{\mathrm{MOR}}$ random effect, and $\Omega = \omega^2$ is the corresponding variance. RSEs refer to the standard deviation estimates, not the variance estimates. ETA shrinkage is reported on the standard deviation and variance scales as provided by NONMEM.

**Supplementary Table S3:** Estimation diagnostics, parameter precision, ETA shrinkage, and selected parameter correlations across four CYP2D6 model specifications.

**A. Estimation diagnostics**

| Diagnostic | AS pooled | AS grouped | Categorical pooled | Categorical grouped |
|---|---|---|---|---|
| Minimization | Successful (with warning) | Successful | Successful | Successful |
| Covariance step | Completed | Completed | Completed | Completed |
| Significant digits | 3.5 | 4.1 | 3.5 | 3.5 |
| Function evaluations | 216 | 1082 | 1205 | 1066 |
| Rounding errors | Not reported | Not reported | Not reported | Not reported |
| Boundary warnings | Not reported | Not reported | Not reported | Not reported |

**B. Variability of the morphine formation fraction and ETA shrinkage**

| Fixed effect / IIV | Group | Ω variance | SE(Ω) | RSE(Ω), % | ETA SD | RSE(SD), % | Shrinkage SD, % | Shrinkage variance, % |
|---|---|---|---|---|---|---|---|---|
| AS pooled | All | 0.334 | 0.014 | 4.2 | 0.578 | 2.1 | 8.7 | 16.6 |
| AS grouped | PM | 5.79 | 3.48 | 60.1 | 2.41 | 30.0 | 80.9 | 96.3 |
| AS grouped | IM | 0.234 | 0.017 | 7.4 | 0.484 | 3.7 | 53.7 | 78.6 |
| AS grouped | NM | 0.119 | 0.0051 | 4.3 | 0.345 | 2.2 | 35.6 | 58.6 |
| AS grouped | UM | 0.326 | 0.065 | 20.0 | 0.571 | 10.0 | 77.0 | 94.7 |
| Categorical pooled | All | 0.185 | 0.0072 | 3.9 | 0.430 | 1.9 | 13.5 | 25.1 |
| Categorical grouped | PM | 0.355 | 0.114 | 32.2 | 0.596 | 16.1 | 82.3 | 96.9 |
| Categorical grouped | IM | 0.289 | 0.025 | 8.6 | 0.538 | 4.3 | 52.8 | 77.7 |
| Categorical grouped | NM | 0.133 | 0.0059 | 4.5 | 0.365 | 2.2 | 34.3 | 56.9 |
| Categorical grouped | UM | 0.110 | 0.024 | 21.6 | 0.332 | 10.8 | 80.0 | 96.0 |

Ω is the estimated variance of the random effect on the logit of the morphine formation fraction ($f_{MOR}$). SE(Ω) and RSE(Ω) refer to the variance scale. ETA SD = $\sqrt{\Omega}$; RSE(SD) was approximated using the delta method as RSE(Ω)/2.

**C. Relative standard errors of selected fixed-effect PK parameters (%)**

| Parameter | AS pooled | AS grouped | Categorical pooled | Categorical grouped |
|---|---|---|---|---|
| Codeine absorption rate | 13.2 | 13.4 | 12.8 | 12.8 |
| Codeine clearance | 28.4 | 40.6 | 110.2 | 44.9 |
| Codeine volume | 28.6 | 40.6 | 110.2 | 45.0 |
| Codeine bioavailability | 28.4 | 40.5 | 110.3 | 45.0 |
| Morphine clearance | 31.3 | 47.2 | 113.4 | 49.4 |
| Morphine volume | 34.9 | 50.2 | 114.6 | 51.3 |

RSE = SE/|estimate| × 100%. Estimates and SEs were extracted from the final NONMEM EXT files. The especially large fixed-effect RSEs in the categorical pooled model should be interpreted together with the parameter correlations in panel D.

**D. Selected fixed-effect parameter correlations**

| Parameter pair | AS pooled | AS grouped | Categorical pooled | Categorical grouped |
|---|---|---|---|---|
| Codeine clearance–volume | 0.996 | 0.998 | ≥0.999 | 0.998 |
| Codeine clearance–bioavailability | 0.998 | 0.999 | ≥0.999 | 0.999 |
| Morphine clearance–volume | 0.886 | 0.946 | 0.990 | 0.949 |
| PM–IM formation parameter | — | — | 0.964 | 0.920 |
| IM–NM formation parameter | — | — | 0.992 | 0.988 |
| NM–UM formation parameter | — | — | 0.959 | 0.981 |

Correlations are estimates of fixed-effect parameter correlations from the NONMEM covariance output (not correlations between individual random effects). Other correlations use matching structural parameters across models. Values are rounded to three decimals; ≥ 0.999 indicates a value printed as 1.00 in the NONMEM listing and does not imply an exact correlation of 1.

AS, CYP2D6 activity score; IIV, interindividual variability; PM, poor metabolizer; IM, intermediate metabolizer; NM, normal metabolizer; UM, ultrarapid metabolizer; RSE, relative standard error; SE, standard error; ETA, individual random effect; PK, pharmacokinetic.

**Supplementary Table S4:** Estimation diagnostics for the exploratory three-group IIV model.

| **Item** | **Reported value** |
|---|---|
| OFV without constant | −2135.3642 |
| Rounding errors | Reported |
| Termination problems | Reported |
| Covariance step | Failed (COV step error) |
| $\theta_1$ Codeine KA | 8.324 |
| $\theta_2$ Codeine CL | 108.1 |
| $\theta_3$ Codeine V | 412.6 |
| $\theta_4$ Codeine bioavailability | 0.8173 |
| $\theta_5$ Morphine CL | 290.1 |
| $\theta_6$ Morphine V | 17.18 |
| $\theta_7$ Reference morphine formation parameter (AS = 2) | 0.1321 |
| $\theta_8$ CYP2D6 genetic effect | 0.9711 |
| $\theta_9$ C6G CL | 8.025 |
| $\theta_{10}$ C6G V | 5.413 |
| $\theta_{11}$ M3G CL | 7.651 |
| $\theta_{12}$ M3G V | 6.911 |
| SD of $\eta_{f_{\mathrm{MOR}}}$, PM + IM | 0.8243 |
| SD of $\eta_{f_{\mathrm{MOR}}}$, NM | 0.3551 |
| SD of $\eta_{f_{\mathrm{MOR}}}$, UM | 0.8360 |
| ETA shrinkage, SD (%) | 47.1; 35.4; 75.9 |
| ETA shrinkage, variance (%) | 72.0; 58.2; 94.2 |

The exploratory model specified three $f_{\mathrm{MOR}}$ random effects: one shared by poor and intermediate metabolizers (PM + IM), one for normal metabolizers (NM), and one for ultrarapid metabolizers (UM). Random-effect standard deviations were obtained from the NONMEM output; the corresponding variances were 0.6794, 0.1261, and 0.6989, respectively. Estimation terminated with numerical errors, and the covariance step failed. Relative standard errors were therefore unavailable.

**Supplementary Table S5:** Model-implied distributions of morphine formation clearance across four independently fitted pharmacokinetic models.

| CYP2D6 fixed effect | IIV structure | Phenotype | Mean (SD) | Median | P5–P95 | CV (%) |
|---|---|---|---|---|---|---|
| Continuous AS | Pooled | PM | 2.70 (1.66) | 2.31 | 0.89–5.90 | 61.4 |
| Continuous AS | Pooled | IM | 6.57 (3.94) | 5.66 | 2.11–14.03 | 59.9 |
| Continuous AS | Pooled | NM | 15.49 (8.14) | 13.91 | 5.40–31.25 | 52.5 |
| Continuous AS | Pooled | UM | 35.42 (14.75) | 33.33 | 15.24–62.96 | 41.6 |
| Continuous AS | Phenotype-specific | PM | 11.68 (20.81) | 2.48 | 0.05–61.38 | 178.2 |
| Continuous AS | Phenotype-specific | IM | 5.22 (2.69) | 4.68 | 2.00–10.35 | 51.5 |
| Continuous AS | Phenotype-specific | NM | 9.55 (3.44) | 9.09 | 4.82–15.82 | 36.0 |
| Continuous AS | Phenotype-specific | UM | 19.91 (9.99) | 17.99 | 7.56–38.69 | 50.2 |
| Categorical phenotype | Pooled | PM | 0.28 (0.13) | 0.252 | 0.12–0.52 | 46.0 |
| Categorical phenotype | Pooled | IM | 4.92 (2.22) | 4.48 | 2.15–9.15 | 45.3 |
| Categorical phenotype | Pooled | NM | 10.18 (4.20) | 9.48 | 4.71–17.91 | 41.3 |
| Categorical phenotype | Pooled | UM | 13.17 (5.24) | 12.34 | 6.25–22.75 | 39.8 |
| Categorical phenotype | Phenotype-specific | PM | 0.26 (0.17) | 0.215 | 0.08–0.58 | 66.3 |
| Categorical phenotype | Phenotype-specific | IM | 4.34 (2.45) | 3.79 | 1.54–8.88 | 56.4 |
| Categorical phenotype | Phenotype-specific | NM | 8.51 (3.08) | 8.04 | 4.37–14.20 | 36.2 |
| Categorical phenotype | Phenotype-specific | UM | 11.00 (3.57) | 10.48 | 6.13–17.54 | 32.5 |

Morphine formation clearance ($CL_{\text{FORM}}$) is expressed in L/h. Each model used its own fitted parameter set and 10,000 simulated individuals per phenotype. CV = 100 × SD/mean. Continuous AS summaries come from the fitted scenarios of the earlier controlled AS simulation, while categorical summaries come from the four-model simulation. No hybrid scenarios are included. These descriptive summaries use the data underlying Figure 1.

**Supplementary Table S6:** Controlled effect of phenotype-specific $f_{\mathrm{MOR}}$ interindividual variability on simulated morphine formation clearance.

**A. Simulated morphine formation clearance (CLFORM)**

| Phenotype / scenario | Mean | SD | Median | P5 | P95 | CV (%) | Sample size |
|---|---|---|---|---|---|---|---|
| PM / A | 0.28 | 0.13 | 0.25 | 0.12 | 0.53 | 46.6 | 10,000 |
| PM / B | 0.30 | 0.20 | 0.25 | 0.09 | 0.68 | 66.0 | 10,000 |
| PM / C | 0.24 | 0.11 | 0.21 | 0.10 | 0.44 | 47.5 | 10,000 |
| PM / D | 0.25 | 0.17 | 0.21 | 0.08 | 0.58 | 65.7 | 10,000 |
| IM / A | 4.92 | 2.19 | 4.52 | 2.19 | 9.02 | 44.5 | 10,000 |
| IM / B | 5.15 | 2.84 | 4.52 | 1.84 | 10.65 | 55.1 | 10,000 |
| IM / C | 4.15 | 1.84 | 3.81 | 1.83 | 7.66 | 44.4 | 10,000 |
| IM / D | 4.34 | 2.41 | 3.80 | 1.57 | 8.92 | 55.6 | 10,000 |
| NM / A | 10.18 | 4.20 | 9.47 | 4.75 | 18.15 | 41.2 | 10,000 |
| NM / B | 10.00 | 3.54 | 9.47 | 5.24 | 16.56 | 35.4 | 10,000 |
| NM / C | 8.72 | 3.69 | 8.07 | 4.07 | 15.48 | 42.4 | 10,000 |
| NM / D | 8.49 | 3.10 | 7.98 | 4.43 | 14.31 | 36.5 | 10,000 |
| UM / A | 13.25 | 5.37 | 12.37 | 6.18 | 23.08 | 40.5 | 10,000 |
| UM / B | 12.89 | 4.16 | 12.35 | 7.27 | 20.66 | 32.3 | 10,000 |
| UM / C | 11.36 | 4.66 | 10.57 | 5.21 | 20.23 | 41.0 | 10,000 |
| UM / D | 10.92 | 3.58 | 10.40 | 6.00 | 17.61 | 32.8 | 10,000 |

*Notes.* CLFORM, morphine formation clearance (L/h); CV, coefficient of variation; P5 and P95, 5th and 95th percentiles; pp, percentage points; PM, poor metabolizer; IM, intermediate metabolizer; NM, normal metabolizer; UM, ultrarapid metabolizer.

Scenario A uses the fitted categorical pooled-IIV model. Scenario B retains all parameters from that model except for the replacement of pooled $f_{\mathrm{MOR}}$ IIV with phenotype-specific IIV. Scenario C retains all parameters from the fitted categorical phenotype-specific-IIV model except for the replacement of phenotype-specific $f_{\mathrm{MOR}}$ IIV with pooled IIV. Scenario D uses the fitted categorical phenotype-specific-IIV model.

All scenarios used the same virtual covariate cohort (10,000 individuals per phenotype), with independent Monte Carlo draws across scenarios. Hybrid scenarios B and C were not re-estimated from clinical data.

**Supplementary Table S6 (continued).** Controlled effect of phenotype-specific $f_{\mathrm{MOR}}$ interindividual variability on simulated morphine formation clearance.

**B. Controlled changes in CLFORM distributions**

| Phenotype / contrast | Mean ratio | SD ratio | Median ratio | P5 ratio | P95 ratio | ΔCV (pp) |
|---|---|---|---|---|---|---|
| PM / A→B | 1.085 | 1.536 | 0.998 | 0.749 | 1.292 | +19.4 |
| PM / C→D | 1.080 | 1.494 | 0.996 | 0.743 | 1.301 | +18.2 |
| IM / A→B | 1.047 | 1.298 | 0.999 | 0.838 | 1.181 | +10.6 |
| IM / C→D | 1.047 | 1.311 | 0.998 | 0.861 | 1.164 | +11.2 |
| NM / A→B | 0.982 | 0.844 | 1.000 | 1.103 | 0.913 | -5.8 |
| NM / C→D | 0.974 | 0.840 | 0.990 | 1.089 | 0.924 | -5.8 |
| UM / A→B | 0.974 | 0.775 | 0.999 | 1.177 | 0.895 | -8.2 |
| UM / C→D | 0.961 | 0.769 | 0.985 | 1.152 | 0.870 | -8.2 |

*Notes.* CLFORM, morphine formation clearance (L/h); CV, coefficient of variation; P5 and P95, 5th and 95th percentiles; pp, percentage points; PM, poor metabolizer; IM, intermediate metabolizer; NM, normal metabolizer; UM, ultrarapid metabolizer.

Scenario A uses the fitted categorical pooled-IIV model. Scenario B retains all parameters from that model except for the replacement of pooled $f_{\mathrm{MOR}}$ IIV with phenotype-specific IIV. Scenario C retains all parameters from the fitted categorical phenotype-specific-IIV model except for the replacement of phenotype-specific $f_{\mathrm{MOR}}$ IIV with pooled IIV. Scenario D uses the fitted categorical phenotype-specific-IIV model.

All scenarios used the same virtual covariate cohort (10,000 individuals per phenotype), with independent Monte Carlo draws across scenarios. Hybrid scenarios B and C were not re-estimated from clinical data.

In Part B, ratios represent the alternative scenario divided by the reference scenario. Changes in CV are expressed as absolute percentage-point differences. Contrasts were calculated from unrounded summary statistics.

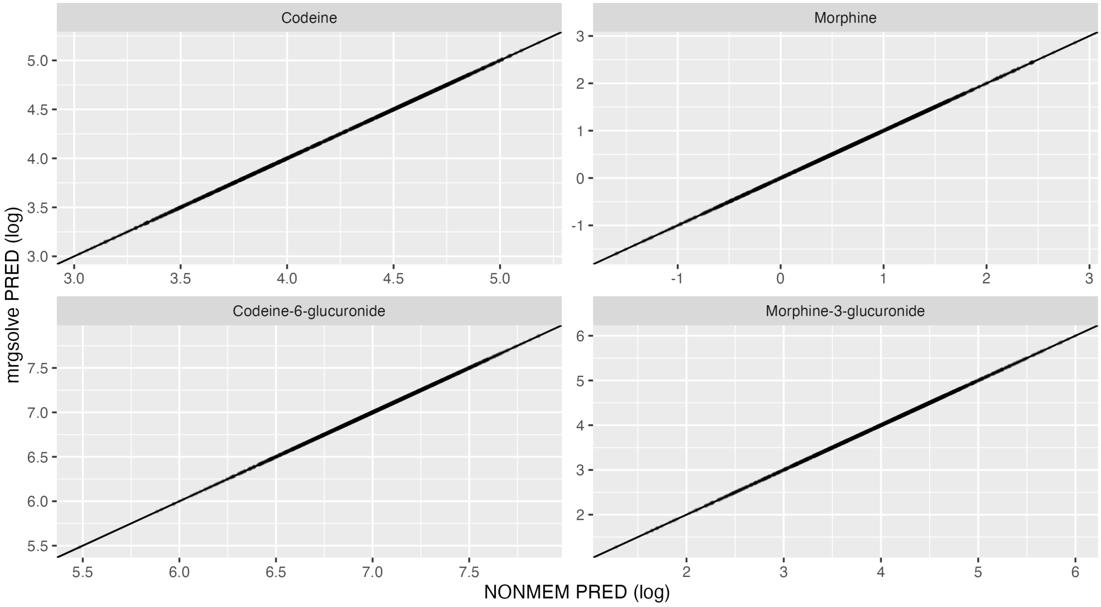


**Supplementary Figure S1:** Deterministic validation of the NONMEM-to-mrgsolve translation of the original published model. Population predictions from NONMEM and mrgsolve are compared on the log-concentration scale for codeine, morphine, codeine-6-glucuronide, and morphine-3-glucuronide. The diagonal line represents the line of identity. Random effects and residual variability were disabled.

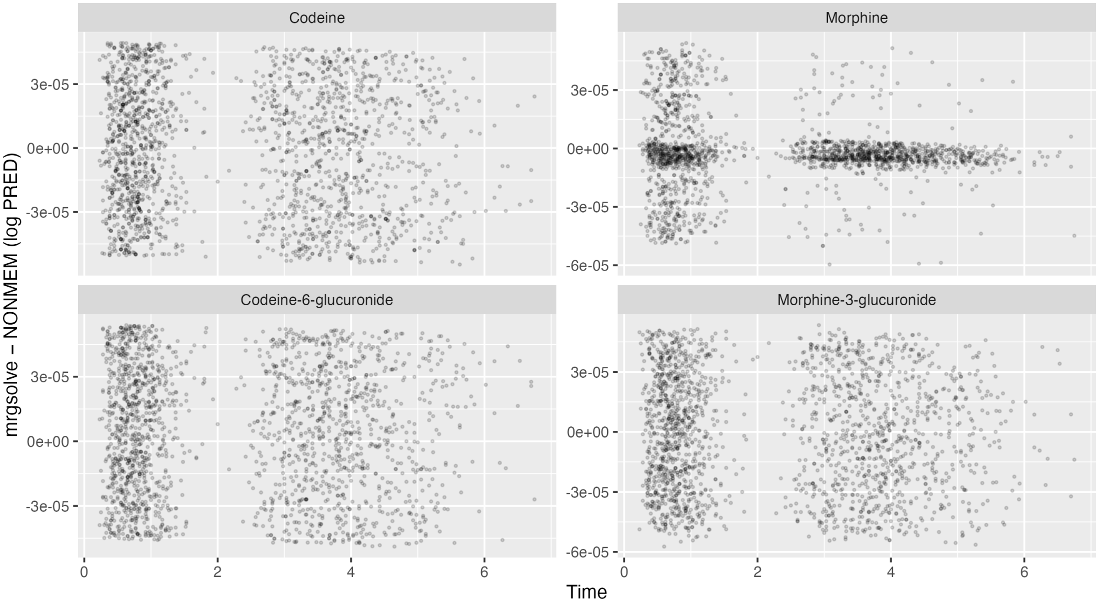


**Supplementary Figure S2:** Differences between mrgsolve and NONMEM population predictions over time for the original published model. Differences were calculated on the log-concentration scale for all four analytes under deterministic conditions. Values close to zero indicate agreement between the two implementations.

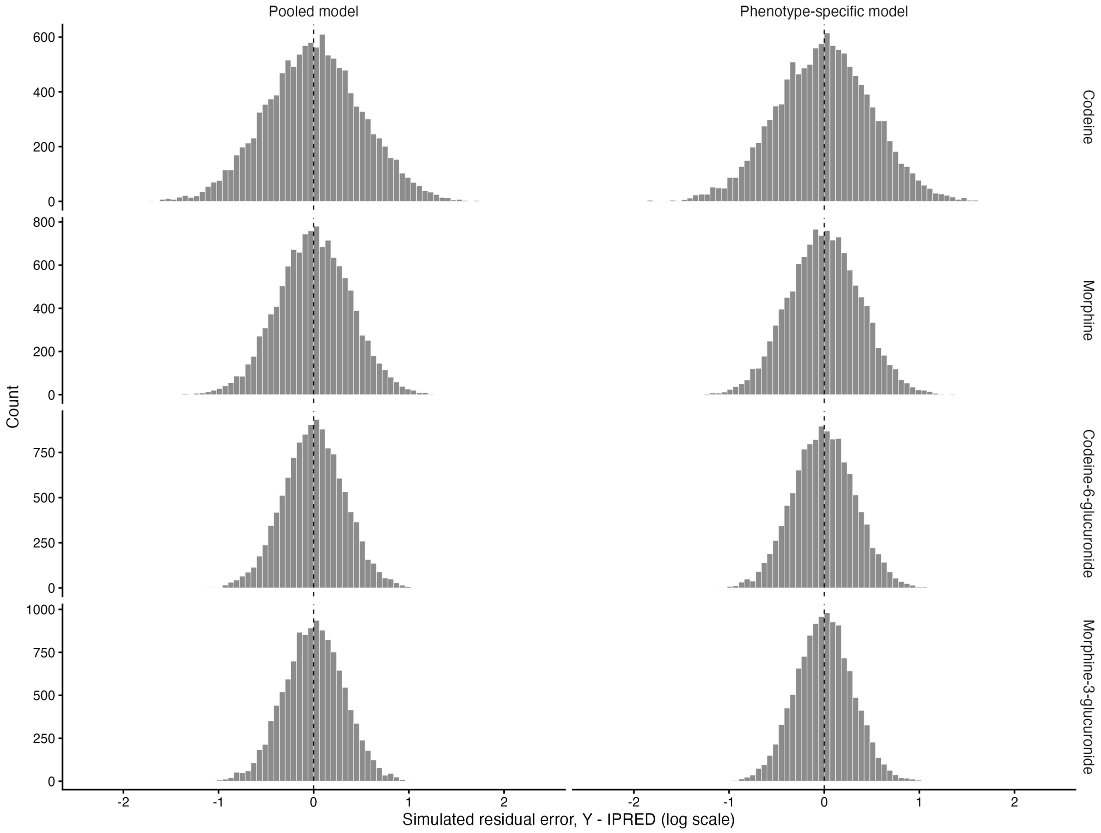


**Supplementary Figure S3:** Stochastic validation of the residual error model under the continuous activity-score parameterization. Histograms show simulated residual errors (Y - IPRED) on the log-concentration scale for codeine, morphine, codeine-6-glucuronide, and morphine-3-glucuronide under the pooled and phenotype-specific models. Each model was evaluated using 3,000 virtual individuals per CYP2D6 phenotype. Dashed vertical lines indicate zero residual error. The same histogram bin width and horizontal axis limits were used for both models.

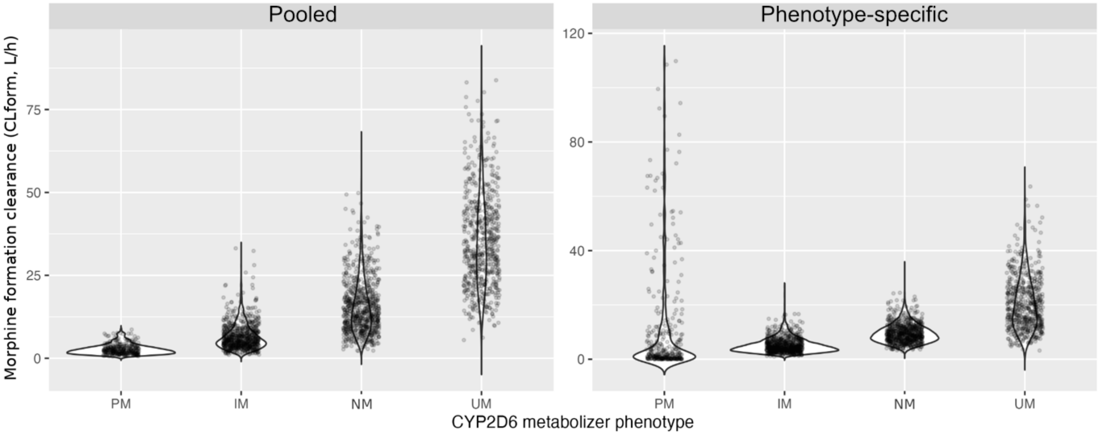


**Supplementary Figure S4:** Violin plots of simulated morphine formation clearance by CYP2D6 phenotype under the continuous activity-score parameterization. Violin plots show the distributions of morphine formation clearance in poor (PM), intermediate (IM), normal (NM), and ultrarapid (UM) metabolizers under the pooled and phenotype-specific IIV models. Points represent a random sample of simulated individuals. The models used their respective estimated parameter sets and the same cohort of 10,000 virtual individuals. The vertical scales differ between panels.

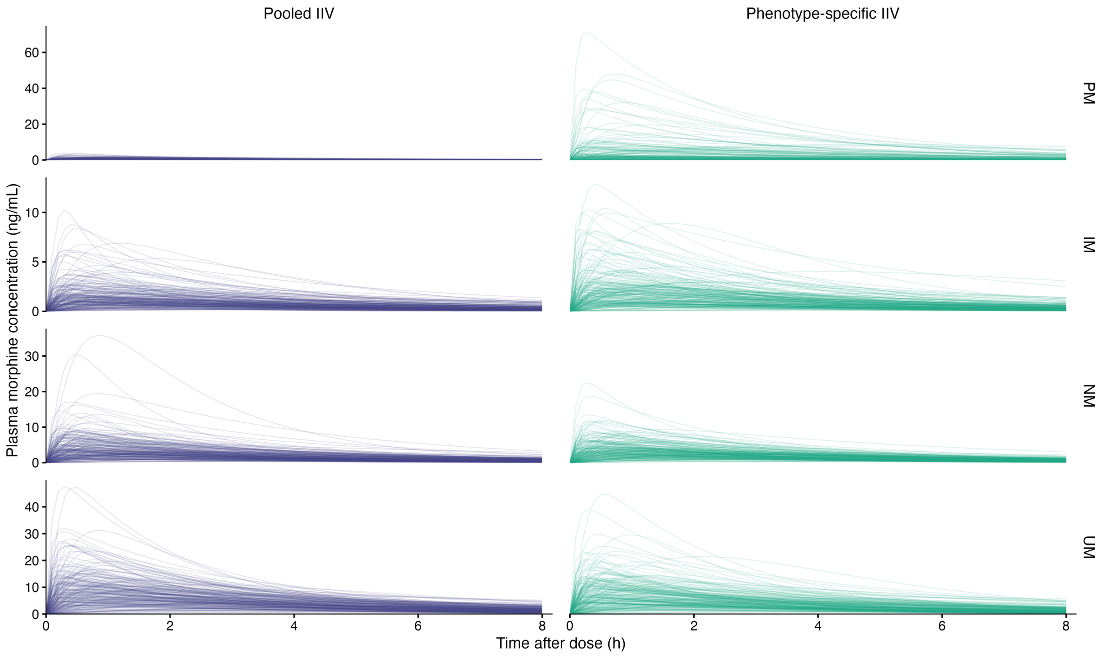


**Supplementary Figure S5:** Individual simulated morphine concentration–time profiles by CYP2D6 phenotype under the continuous activity-score parameterization. Profiles are shown for 200 randomly selected individuals per phenotype and model following a single oral dose of 60 mg codeine. Individuals were selected from the simulation cohort used for Figure 3, with the same resampled covariates but independently drawn random effects between models. Blue and green lines represent the pooled and phenotype-specific IIV models, respectively. Concentration axes differ between phenotypes but are identical between models within each phenotype. Simulated concentrations do not include residual error.
